# The impact of paediatric epilepsy and co-occurring neurodevelopmental disorders on functional brain networks in wake and sleep

**DOI:** 10.1101/2023.03.15.530959

**Authors:** Leandro Junges, Daniel Galvis, Alice Winsor, Grace Treadwell, Caroline Richards, Stefano Seri, Samuel Johnson, John R. Terry, Andrew P. Bagshaw

## Abstract

Epilepsy is one of the most common neurological disorders in children. Diagnosing epilepsy in children can be very challenging, especially as it often coexists with neurodevelopmental conditions like autism and ADHD. Functional brain networks obtained from neuroimaging and electrophysiological data in wakefulness and sleep have been shown to contain signatures of neurological disorders, and can potentially support the diagnosis and management of co-occurring neurodevelopmental conditions. In this work, we use electroencephalography (EEG) recordings from children, in restful wakefulness and sleep, to extract functional connectivity networks in different frequency bands. We explore the relationship of these networks with epilepsy diagnosis and with measures of neurodevelopmental traits, obtained from questionnaires used as screening tools for autism and ADHD. We explore differences in network markers between children with and without epilepsy in wake and sleep, and quantify the correlation between such markers and measures of neurodevelopmental traits. Our findings highlight the importance of considering the interplay between epilepsy and neurodevelopmental traits when exploring network markers of epilepsy.

## Introduction

Epilepsy is estimated to impact nearly 10.5 million children worldwide^1^. In addition to the personal, social, and economic impact of epilepsy on children and their families, seizures have been shown to be detrimental to brain development^2^, potentially leading to cognitive dysfunction, and the condition is often associated with lifelong disabilities and poor quality of life^3,4^. Therefore, early and accurate diagnosis of epilepsy is paramount. Unfortunately, epilepsy diagnosis can be very challenging. The rate of epilepsy misdiagnosis is estimated to be near 20% generally^5^ and, due to a wide range of non-epileptic paroxysmal disorders and co-occurrences affecting children^6^, misdiagnosis in children is believed to be even greater than for adults^7^.

Diagnosis and management of neurological and neurodevelopmental conditions are made more challenging when they coexist. This is frequently the case with epilepsy, where its prevalence in children with Autism Spectrum Disorder and ADHD is 20% and 15%, respectively, which is significantly higher than in neurotypical children (∼1%)^8^. The complex relationship between epilepsy and co-occurring neurodevelopmental conditions remains an important open question, the resolution of which could improve clinical outcomes and provide optimal and individualised care.

Epilepsy is increasingly conceptualised as a condition of aberrant brain networks^9,10^. Scalp electroencephalography (EEG) is one of the most widespread methods used to quantify these networks. Functional networks obtained from scalp EEG have shown fundamental differences between people with epilepsy and healthy controls, for both adults^11,12,13^ and children^14,15^. Neurodevelopmental conditions, such as autism and ADHD, have also been investigated using the framework of network science^16^, although these methods are less well stablished in this context. Moreover, very few studies have investigated the joint effect of epilepsy and co-occurring neurodevelopmental conditions on functional brain networks^17^. This is necessary to understand network signatures that are specific either to epilepsy, or to neurodevelopmental conditions, rather than being sensitive to their co-occurrences. Network markers of epilepsy may be influenced by the presence of neurodevelopmental traits, potentially leading to erroneous interpretations of the relationship between these markers and seizure propensity.

Another important factor when exploring network markers of co-occurring neurological and neurodevelopmental conditions is the influence of sleep. A growing number of studies support the association between poor sleep and both epilepsy and neurodevelopmental conditions ^18,19,20,21,22^. This relationship tends to be bidirectional, where sleep disruption can increase seizure propensity and presentation of neurodevelopmental conditions, which can in turn result in poor sleep^23^. At the same time, graph metric analysis has shown that several aspects of sleep, such as the wake-sleep transition itself as well as sleep deprivation, are associated with connectivity changes in functional brain networks^24,25,26,27,28^. Markers of epilepsy might also be influenced by stages of awareness (wake and sleep), given the known changes to the propensity for epileptic discharges with sleep stage^29,30^.

In this work we explore the combined effects of epilepsy and neurodevelopmental traits on functional connectivity networks obtained from EEG recordings from children in waking restfulness and sleep. We identify differences in functional connectivity between subjects with and without epilepsy, which are consistent across frequency bands. We also show that such differences are less pronounced during sleep. Finally, we quantify the correlation between neurodevelopmental traits and network measures, identifying similar effects as seen for epilepsy. These results highlight the importance of considering the co-occurrence of neurodevelopmental traits when a graph metric approach is implemented in this context.

## Methods

### Data acquisition and participants

The data used in this study were acquired at Birmingham Children’s Hospital and Worcestershire Royal Hospital. Written informed consent and assent were obtained from parents and children and the study received NHS ethical approval from the Northwest - Preston Research Ethics Committee (REC reference 19/NW/0337). EEG recordings were collected in “nap sleep EEG” sessions (i.e., recordings taken during a short period where the child falls asleep) from children suspected of having epilepsy, as part of the diagnostic process. Sixty-two recordings, collected between September 2019 and December 2021, were retrieved. EEG data were acquired from 19 electrodes positioned according to the 10-20 system and sampled at 512Hz. In some participants, melatonin or mild sleep deprivation were used to encourage sleep, according to clinical protocols. Families also completed the Social Communication Questionnaire (SCQ)^31^ and the Conners’ 3AI Questionnaires^32^, which are standard tools to describe autism and ADHD characteristics, respectively. All questionnaires were evaluated by experienced psychologists (AW and CR) to provide continuous indices associated with autism/ADHD traits. Raw scales of the SCQ and Conners’ questionnaires can have values in the ranges of [0,40] and [0,20], respectively.

In order to define a quantity that represented overall neurodevelopmental traits (*NT*), we combined SCQ and Conners’ raw scores as:

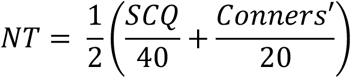

With this definition, *NT* ranges in [0,1], where 0 means a null score in both tests while 1 means maximum scores in both tests. This index allows us to quantify the overall level of neurodevelopmental traits in a single dimension^33^. It is important to clarify here that *NT* should not be interpreted as a detailed quantification of autism and ADHD diagnosis. These conditions have complex diagnostic pathways, which go beyond the interpretation of these questionnaires. However, despite its limitations, these questionnaires (and therefore *NT*) constitute an accessible and informative marker for the characteristics associated with these conditions.

### EEG Analysis

EEG annotation was performed by two experienced electrophysiologists (Neuronostics Ltd). For each participant, electrophysiologists were provided with the complete EEG recording from the nap sleep session (recording duration between 00:28:00 and 03:48:49 [hh:mm:ss]) and asked to identify the cleanest and most “uneventful” 30-second long EEG segments (avoiding major artifacts or clear epileptiform activity) in wakefulness, and sleep stages N1 - N3, when available. Sleep stages were defined according to AASM guidelines^34^. Very few epochs were identified in sleep stage N3, so those were not considered in this analysis. Electrophysiologists were blind to epilepsy diagnosis and to any metadata associated with neurodevelopmental traits.

### Final Cohort

From the original 62 participants, 34 had at least one EEG epoch identified and complete metadata available (age, sex, epilepsy diagnosis, SCQ score and Conners’ score) and were included in this study. These participants were aged between 4 and 15 years old (median 9 y) and included 13 females and 21 males. 24 participants were diagnosed with epilepsy (11 focal, 7 generalised, 4 Rolandic, and 2 Encephalopathy) while 10 were not. These groups will be referred to as “epilepsy” and “controls”, respectively. Raw scores for the SCQ ranged between 0 and 27 (median 9), while Conners’ raw score ranged between 0 and 20 (median 9.5). See Supplemental Material for detailed metadata.

### Functional Networks

We derived weighted undirected functional networks from each EEG epoch using the phase locking factor (PLF). To do this, we first downsampled the data to 256 Hz and band-pass filtered between the desired frequencies. A 4^th^ order Butterworth filter was used with forward and backward filtering to minimise phase distortions. Functional networks were calculated in five frequency bands: delta (1Hz-4Hz), theta (4Hz-7Hz), alpha (7Hz-13Hz) and beta (13Hz-30Hz), as well as low alpha (6Hz-9Hz). Low alpha was used as networks calculated in this frequency band in adults have shown different properties in healthy individuals and those with generalized epilepsy^11^.

For a pair of signals *k* and *l*, the *PLF*_*kl*_ is given by 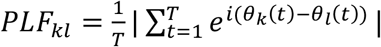, where *T* is the number of equally-spaced time samples in an epoch and *θ*_*k*_ is the phase of the Hilbert transform of signal *k*. We also calculated the time-averaged lag, 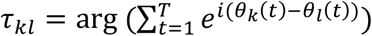. Only nonzero time lags (|*τ*_*kl*_| > 0) were considered to avoid spurious connections due to volume conduction. We then computed 99 surrogate epochs from each of the EEG signals using a univariate iterated amplitude adjusted Fourier transform (iAAFT). Functional networks were then calculated for the EEG epochs and for the surrogates. For each epoch, we rejected connections that did not exceed a 95% significance level compared to the same connection weights computed from the surrogates calculated for that epoch. This method results in a weighted, undirected network *a*_*kl*_, which we used to calculate graph metrics. Details about the methods used to calculate the network mean degree (MD), degree standard deviation (DStd), average local clustering coefficient (ALCC) and global efficiency (GE) can be found in the Supplemental Material.

### Statistical Analysis

We explored the weighted mean degree of different classes (controls and epilepsy types) using boxplots (see Fig. 1), where the median (red line), 25^th^ - 75^th^ percentiles (blue box), non-outlier extremes (black dashed lines) and outliers (red crosses) of the distributions are presented. Effect size was quantified using the rank-biserial correlation^35^ (|*r*| ∈ [0,1], where 0 means no rank correlation and 1 means perfect separation between groups), and significance was calculated using the Wilcoxon rank-sum and Kruskal-Wallis tests. To further quantify the differences between classes, receiver operating characteristic (ROC) curves were calculated for all frequency bands. The area under the ROC curve (AUC) was calculated and uncertainty (error bars) was quantified using a leave-one-out approach. To quantify the relationship between mean degree and neurodevelopmental traits (continuous index), we used the nonparametric Spearman rank correlation measure.

**Figure 1:**
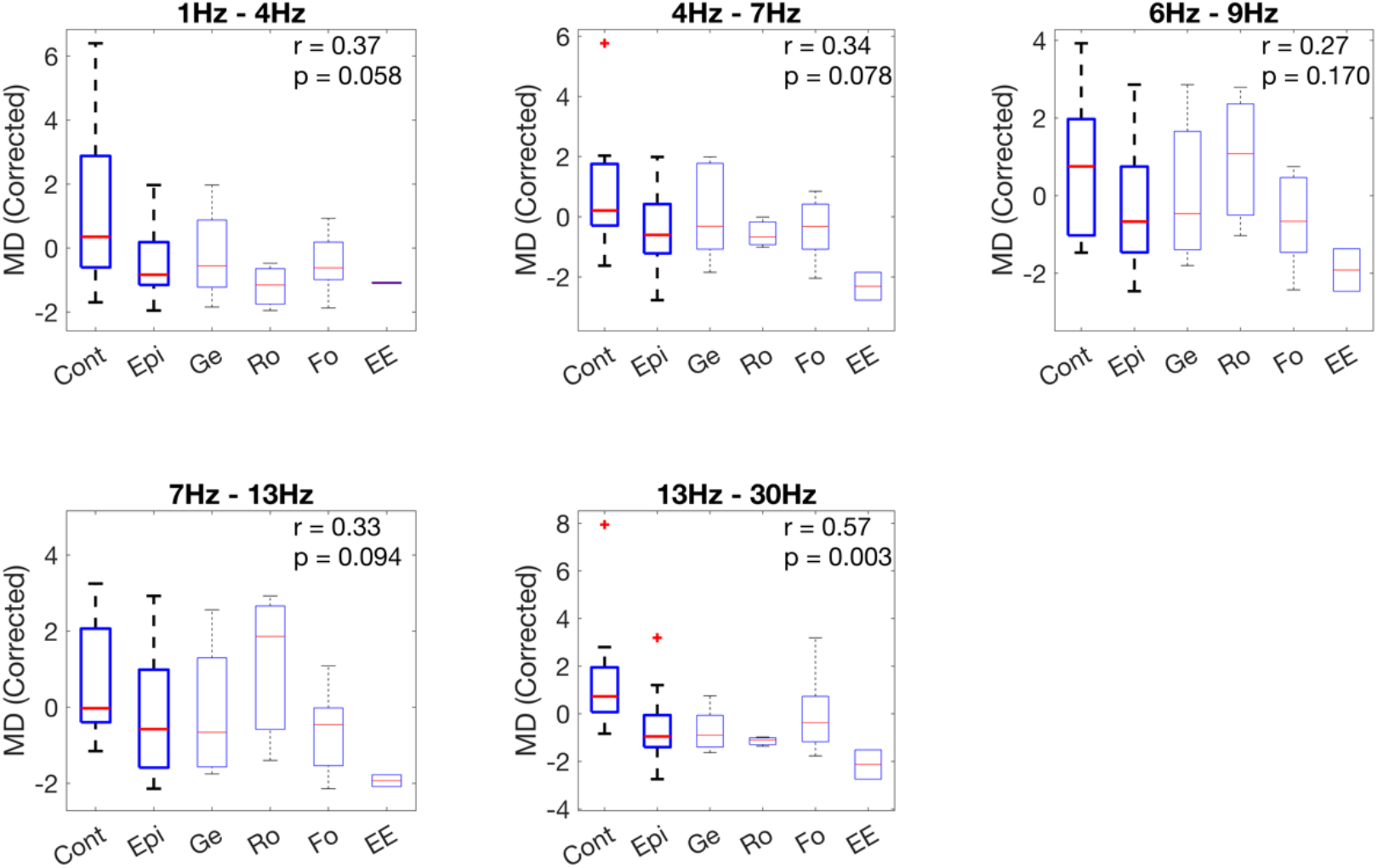
Summary statistics of the functional connectivity networks’ mean degree (corrected for sex), calculated for different frequency bands and using wake epochs. The first two boxes in each plot (“Cont” and “Epi”) indicate the comparison between subjects without and with epilepsy, respectively. Subsequent boxes show the breakdown of different epilepsy types (Ge: generalised, Ro: Rolandic, Fo: focal, and EE: encephalopathy). The rank-biserial correlation and p-value (two-tailed Wilcoxon rank sum test, uncorrected for multiple comparisons) for the difference between “Cont” and “Epi” are also shown.

When comparing controls and epilepsy groups, the age distributions were not significantly different (p-value: 0.79), however there was a clear sex imbalance (controls: 60% female, epilepsy: 29% female), so we corrected the marker values for sex in all comparisons presented below by subtracting the mean over the respective sex. Regarding the correction for confounding factors for the *NT* index, no significant differences were observed between epilepsy and controls, or between males and females. Also, no significant correlation was observed between the *NT* index and age. Nevertheless, to avoid cumulative effects of potential confounding factors, when considering relationships between *NT* and mean degree, we corrected this network marker for age, sex, and epilepsy diagnosis using linear regression.

## Results

### Mean degree is smaller in epilepsy compared to controls

The mean node degree calculated using functional connectivity networks obtained from EEG epochs during wakefulness is presented in Fig. 1. Each plot describes the summary statistics of the mean degree distribution for the different frequency bands of interest. The first two boxes in each plot indicate the mean degree distribution for control and epilepsy subjects, respectively. The subsequent boxes, in faded colours, indicate results for the sub-groups of epilepsy types (Ge: generalised, Ro: Rolandic, Fo: focal, and EE: encephalopathy). For all frequency bands, the median mean degree calculated for subjects with epilepsy was lower than for controls. This result was not only consistent across frequency bands, but also held when controls were compared with most epilepsy types individually. Rolandic epilepsy presented mean degree values similar to controls in the low-alpha and alpha bands. However, it is important to note that this group consisted of only 4 subjects, so any comparison for this group in isolation has to be considered carefully. The rank-biserial correlations presented in each plot indicate that the difference between the mean degree for controls and subjects with epilepsy was clearer in the beta band. We also quantified the differences between controls and epilepsy in degree standard deviation (DStd), average weighted clustering coefficient (AWCC), and global efficiency (GE). We observed trends that were consistent over all frequency bands (elevated DStd and GE for controls and elevated AWCC for children with epilepsy). However, the effect sizes were small (see Fig. S1 in the Supplemental Material) and these metrics were not considered further.

To further quantify the differences between the mean degree for controls and epilepsy, and to estimate its classification power as a marker, we calculated the receiver operating characteristic (ROC) curve, presented in Fig. 2. The area under the ROC curve (AUC) varied between 0.66 and 0.84, depending on the frequency band used to calculate the networks, reflecting the consistent difference observed in the mean degree for controls and epilepsy in Fig. 1.

**Figure 2:**
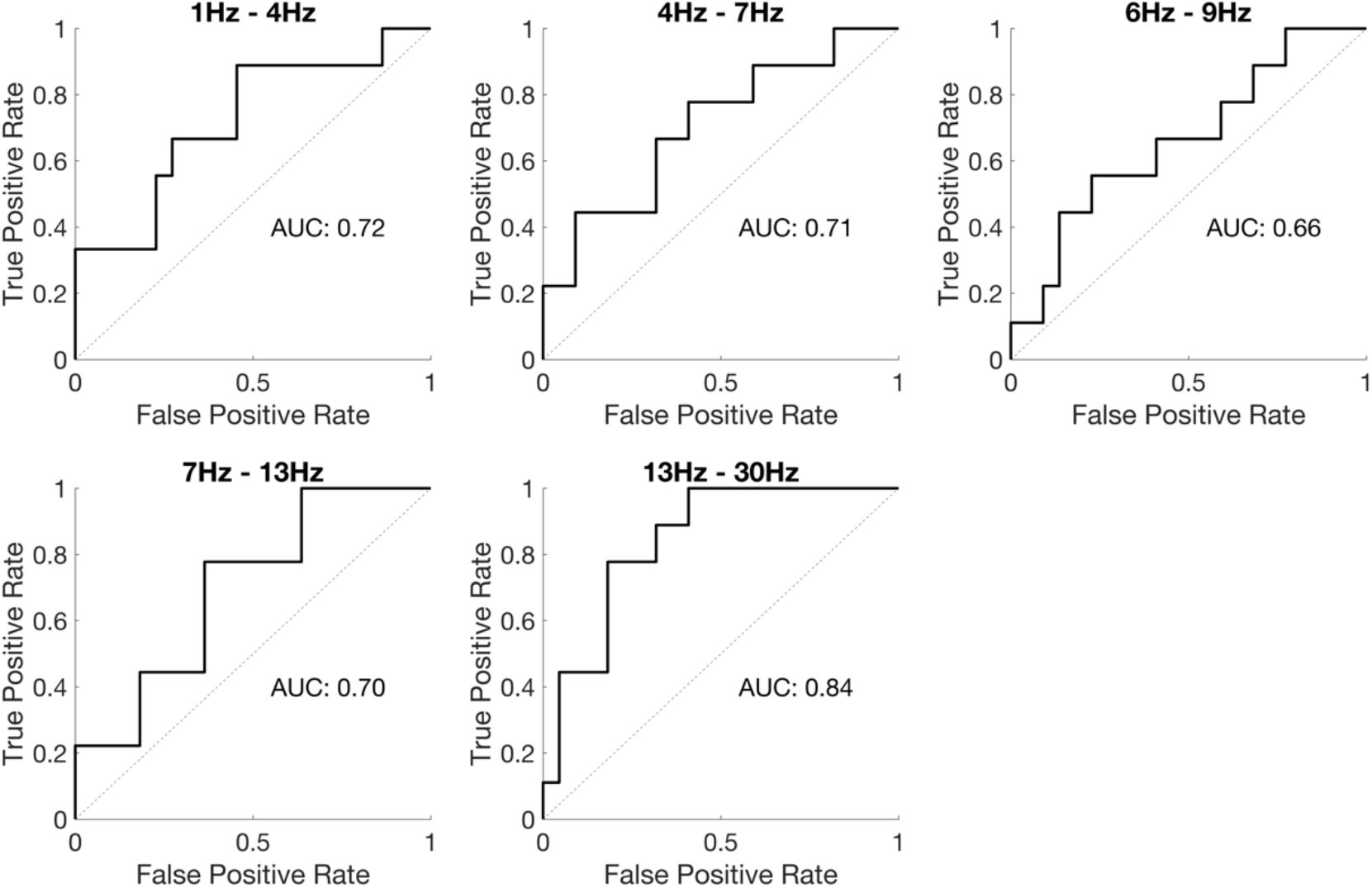
Receiver operating characteristic (ROC) curve calculated using the mean degree to classify subjects without and with epilepsy (wake epochs).

### Differences in mean degree are smaller in sleep compared to wakefulness

As sleep has been shown to be an important factor impacting seizure susceptibility in different types of epilepsy ^23,18^, one important question is how it impacts functional brain networks of children with epilepsy. To answer this question, we calculated differences in mean degree between controls and children with epilepsy for epochs obtained from sleep stages N1 and N2. Following the calculation of the area under the ROC curve for epochs obtained from wakefulness, presented in Fig. 2, we used the AUC to quantify the differences between mean degree for controls and children with epilepsy in sleep (Fig. 3). As subjects transition from wakefulness into sleep (N1 and N2), the differences in mean degree between cases and controls decrease, as evidenced by the decrease in the AUC from wake to N1 and N2 in Fig. 3. For the delta, theta and beta bands, significant differences were observed between wake and N1/N2, while no significant differences were observed between N1 and N2. In the low alpha and alpha bands, no significant differences were observed between wake and N1, while both stages have significantly different AUC than N2 (Kruskal-Wallis test, Bonferroni correction for multiple comparisons).

**Figure 3:**
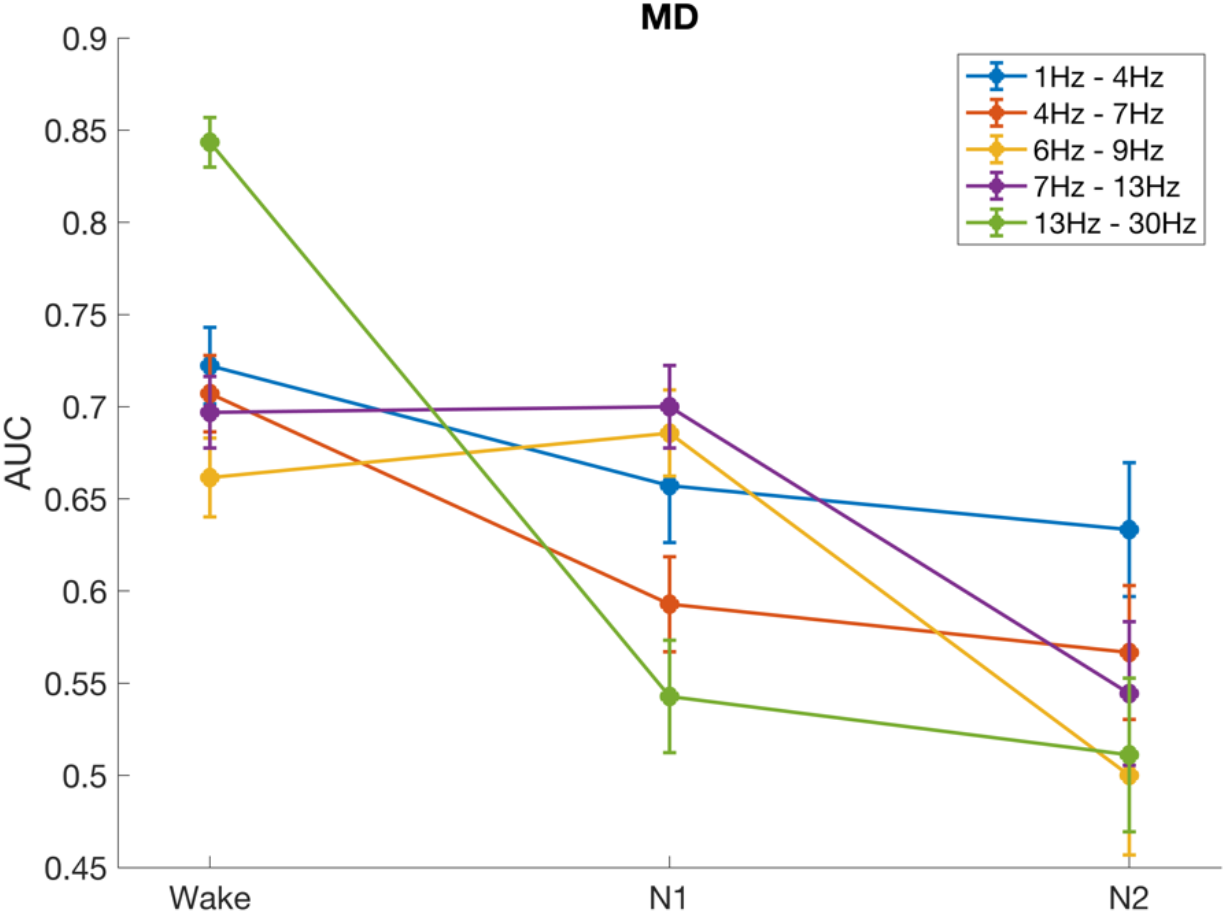
Area under the ROC curve for the mean degree, calculated in different stages of awareness (wake, N1 and N2).

### Neurodevelopmental traits correlate with decrease in mean degree

The effect of autism and ADHD traits on functional brain networks is explored in Fig. 4. In this figure, the network mean degree (corrected for age, sex and epilepsy diagnosis), calculated for wake epochs, was plotted as a function of the neurodevelopmental trait index (see Methods), for all frequency bands. Figure 4 shows a negative correlation between neurodevelopmental traits and mean degree, for all frequency bands. The correlation is clearer for higher frequencies, and remains significant when corrected for multiple comparisons in the low alpha, alpha and beta bands. It is important to note that the mean degree values here are corrected for epilepsy diagnosis (see Methods), so the correlation between mean degree and neurodevelopmental trait index is independent of epilepsy diagnosis. When we consider N1 and N2 epochs, the correlation was generally less clear but followed a similar trend (see Figs. S2 and S3 in the Supplemental Material).

**Figure 4:**
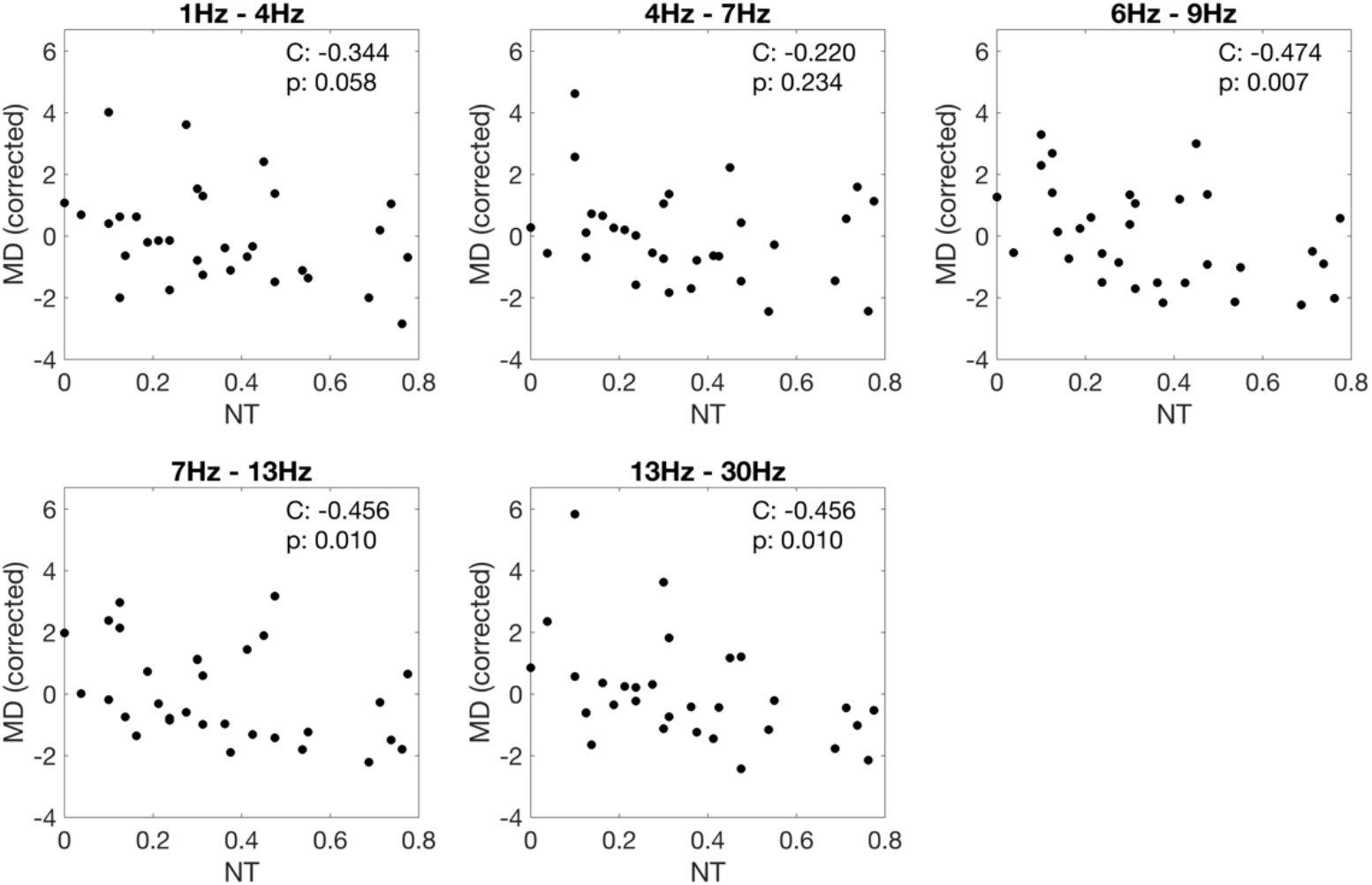
Spearman correlation and p-value (C and p) between neurodevelopmental trait index (*NT*) and mean degree (MD) corrected for age, sex, and epilepsy diagnosis. MD calculated using wake epochs.

The influence of the neurodevelopmental trait index (*NT*) on the mean degree affects the classification of controls and epilepsy subjects using this marker. Fig. 5 (left) shows *NT* and mean degree (corrected for sex), calculated for controls and epilepsy in the low alpha band (which had the highest correlation with *NT*). For the MD threshold of maximum balanced accuracy (dashed line), some subjects were misclassified (blue dots below the dashed line, and red dots above it). When we analysed the *NT* of the misclassified subjects (Fig. 5 - right), we noticed that controls misclassified as epilepsy have a larger median *NT* than controls correctly classified. The opposite effect was seen for epilepsy. The number of misclassified subjects was small, but the trend was clear and consistent across all frequency bands (see Fig. S4 in the Supplemental Material). When estimating the classification power of MD through the calculation of the AUC, if instead of only correcting this marker for sex imbalance (as in Fig. 3) we also correct it for *NT*, the AUC generally improves, especially for wake epochs, as shown in Fig. 6.

**Figure 5:**
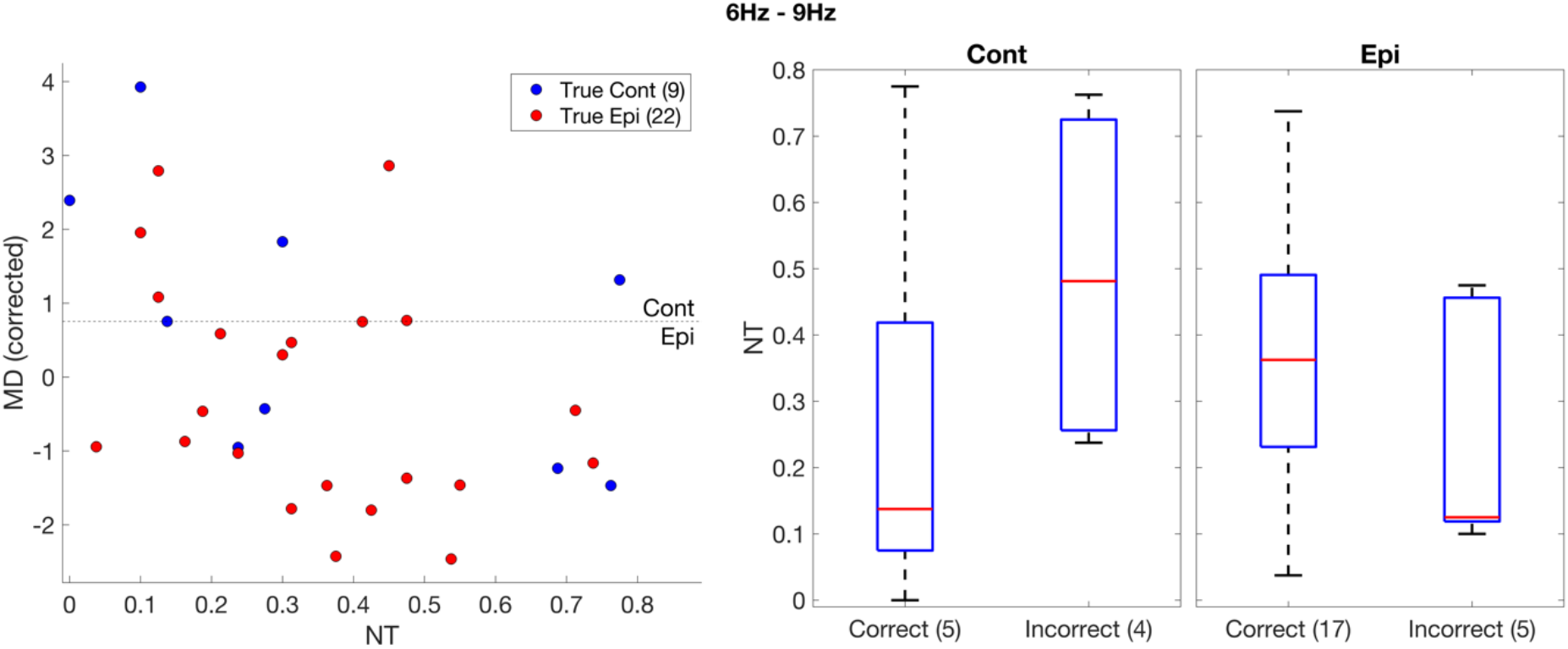
(Left) mean degree corrected for age, calculated using wake epochs, as a function of neurodevelopmental traits. Cont (Epi) are shown in blue (red). The dashed line represents the threshold of optimal balanced accuracy for the separation between Cont and Epi. **(Right)** Comparison between neurodevelopmental traits of subjects classified correctly (Cont > threshold / Epi < threshold) and incorrectly (Cont < threshold / Epi > threshold). Shown here only for low alpha band (6Hz – 9Hz). See Supplemental Material for the same calculation in other frequency bands.

**Figure 6:**
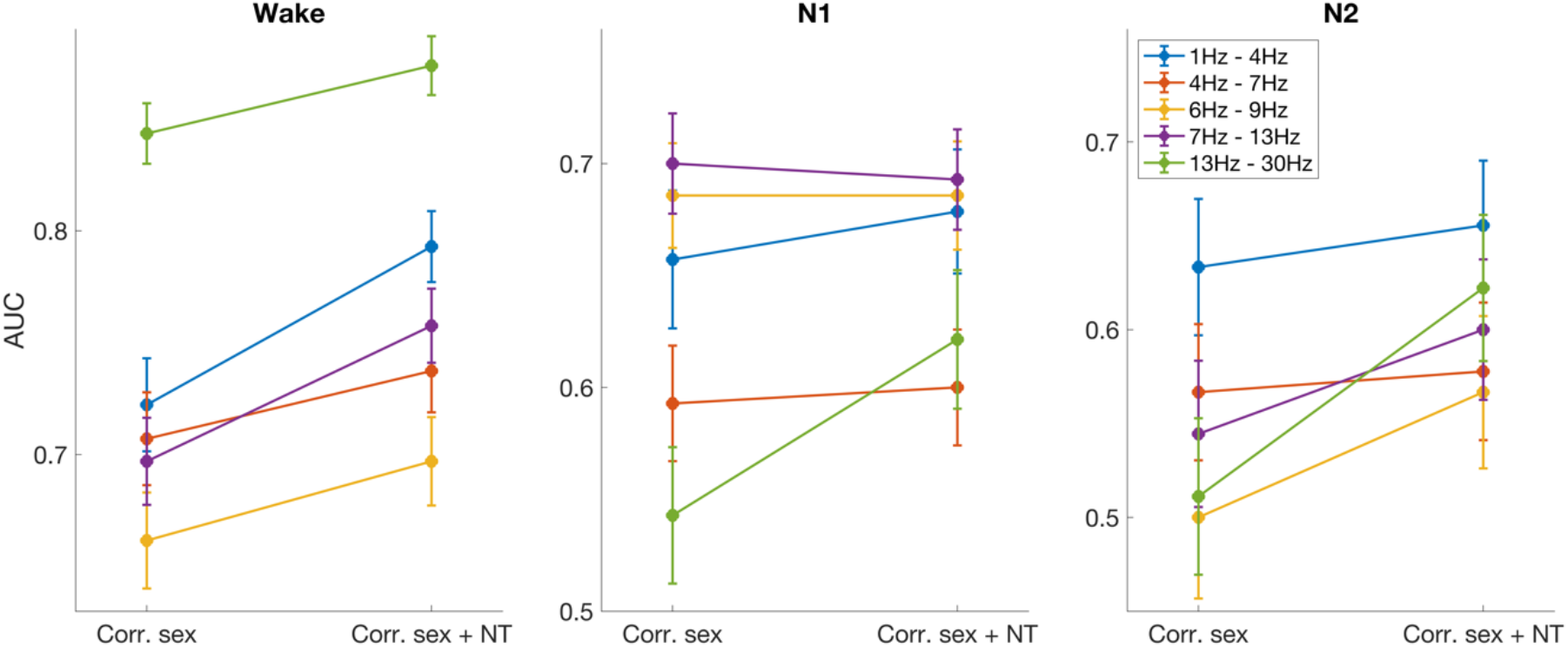
Comparison between AUC calculated using MD only corrected for sex (as in Fig. 3) and corrected for sex and neurodevelopmental traits index (*NT*).

## Discussion

In this work, we investigated how paediatric epilepsy and co-occurring traits of neurodevelopmental conditions impact functional brain networks obtained from EEG in wakeful rest and sleep. We showed that, for networks obtained from wake resting-state epochs, epilepsy diagnosis correlates with a decreased mean degree within different frequency bands, with this effect being most apparent in the beta band. For epochs obtained in sleep stages N1 and N2, this effect is generally less pronounced. We have also shown that a marker associated with autism and ADHD characteristics (*NT*) has a negative correlation with mean degree, which is consistent across frequency bands and stages of awareness. We also quantified how neurodevelopmental traits can influence the classification power of mean degree when separating controls and epilepsy subjects. We showed that children without epilepsy and with high *NT* have a higher risk of being misclassified than those with low *NT*. Conversely, children with epilepsy with low *NT* might have a higher risk of being classified as not having epilepsy if the influence of *NT* is not accounted for when identifying optimal classification thresholds.

Functional networks extracted from resting-state EEG have been studied in the context of epilepsy previously, and various markers have been explored. Chowdhury et al.^11^ compared functional networks from adult controls and adults diagnosed with idiopathic generalised epilepsy. They showed that, in the low-alpha band, network mean degree and degree variance are elevated in epilepsy, while clustering coefficient is lower in epilepsy. These results differ from what has been observed in this work. However, it is important to point out that changes in the pre-processing and calculation of functional networks can have a significant effect on network markers, as can type of epilepsy, so comparisons across different studies need to be interpreted carefully. Potential differences between the effects of epilepsy on network markers in children and adults can result from the intricate influence of brain maturation in the paediatric brain. Resting-state functional EEG networks have been shown to present complex band-specific changes during the maturation period (e.g., positive correlation between network segregation and age in the upper alpha band)^36^. These results evidence the importance of considering the influence of brain maturation in the study of epileptogenic brain networks in children. The effects of age were accounted for in the present study, but comparisons were made considering a relatively broad age range (4 to 15 years old). Further studies with larger sample sizes, clustering participants in narrower age ranges, are needed to clarify the influence of brain maturation on EEG networks in the context of epilepsy and neurodevelopmental disorders.

The results described above, observed in networks derived from wakeful rest, were also consistent with those from epochs from sleep stages N1 and N2, however the effect size was generally smaller during sleep. This result is interesting since NREM sleep has been shown to activate interictal epileptiform discharges (IED) in many types of epilepsies^37^, which actually underpins the use of nap studies to support epilepsy diagnosis. However, it is important to notice that smaller control-epilepsy differences for markers in sleep than in wake does not imply that ictal or interictal activity should be less frequent in sleep. The relationship between IEDs and seizure susceptibility is still unclear, with some works suggesting that IEDs can have anti-seizure effects, depending on the underlying physiological mechanisms leading to seizures^38,39^. In this scenario, states where IEDs are more frequent could lead to network representations with features associated to low ictogenicity. The detailed relationship between IEDs and network markers would require long wake and sleep recordings, rich in IEDs, and is beyond the scope of this work.

The influence of neurodevelopmental conditions, like autism and ADHD, on functional networks extracted from EEG data is still an open question. Evidence suggests that autism is characterised by long-range underconnectivity^40^, but this has been challenged and the diversity in methodology makes it difficult to evaluate and compare across studies^41^. In this study we have shown that network mean degree presents a negative correlation with the neurodevelopmental trait index *NT* (autism and ADHD characteristics). This relationship does not comprehensively describe the effect of autism and/or ADHD on functional brain networks, but it shows how the traits associated with these conditions can influence network-based biomarkers and, therefore, their potential clinical value. The trend observed in the relationship between mean degree and *NT* is also observed for the SCQ and Conners’ raw scores separately (data not shown). In order to disentangle the influences of autism and ADHD on network markers, future studies should extend the analysis presented here by considering cases with confirmed clinical diagnoses of these conditions, and focus on the main characteristics that differentiate their classification.

Most studies that explore network markers of epilepsy from EEG recordings tend to exclude subjects with co-occurring conditions from the analysis, especially neurodevelopmental conditions. However, it is often unclear how and to what extent subjects have been tested, especially when sub-clinical traits of neurodevelopmental conditions are considered. The results presented in this work show that ignoring this information can lead to skewed model calibration and inaccurate classification, especially for children with high *NT*. Such inaccuracies could lead to even longer diagnostic delays, misdiagnosis, and inappropriate treatment strategies.

Some limitations of this work need to be considered when interpreting the results presented above. Our analysis was implemented considering a relatively small number of subjects, especially in the control group. Additionally, autism and ADHD traits was not different between the control and epilepsy groups. Previous works suggest that both conditions have a higher prevalence in epilepsy than in typically developing children^8^, indicating that the data used in this study might not be representative of the general population. However, it is important to point out that the “control” group in this work represents children suspected of having epilepsy who had a differential diagnosis. To the best of our knowledge, the expected prevalence of autism and/or ADHD in such a group is unknown. Future works should also focus on stratifying the analyses above in different epilepsy types, presenting a detailed quantification of the influence of each epilepsy syndrome in network markers and their interrelationship with co-occurring conditions.

## Acknowledgements

L.J. acknowledges support from The Waterloo Foundation via a Child Development Fund Research Grant (grant ref. no. 1970-4687). A.W., C.R., and A.P.B. acknowledge support from The Waterloo Foundation (grant ref no. 1970/3346). L.J., D.G. and J.R.T. acknowledge support from the University of Birmingham Dynamic Investment Fund. J.R.T. acknowledges support from the EPSRC (grant ref. no. EP/T027703/1). S.J. acknowledges support from the Alan Turing Institute and the EPSRC (grant ref. EP/N510129/1).

## Notes

### Competing Interest Statement

John Terry is co-founder and managing director of Neuronostics Ltd.

