## Supplemental Material for "The impact of paediatric epilepsy and co-occurring neurodevelopmental disorders on functional brain networks in wake and sleep"

### Network markers

**Mean Degree (MD):** The weighted degree for a node $i$ is given by $\sum_{j=1}^{N} a_{ij}$ , and the weighted mean degree is given by $\frac{1}{N}\sum_{i=1}^{N}\sum_{j=1}^{N} a_{ij}$.

**Degree standard deviation (DStd):** The weighted-degree standard deviation is given by $\sqrt{\frac{1}{N-1}\sum_{i=1}^{N}|D_i - MD|^2}$, where $D_i = \sum_{j=1}^{N} a_{ij}$ is the weighted degree of node $i$.

**Average local clustering coefficient (ALCC):** The weighted local clustering coefficient for a node $i$ is given by:

$$C_i = \frac{1}{k_i(k_i - 1)}\sum_{j=1}^{N}\sum_{k=1}^{N}\left(a_{ij}a_{jk}a_{ki}\right)^{\frac{1}{3}},$$

where $C_i$ is defined to be 0 if $k_i \leq 1$. $k_i$ is the unweighted degree of node $i$, given by $k_i = |\{a_{ij}|\ a_{ij} > 0\}|$ (the size of the set containing all nonzero edges between node $i$ and the other nodes). The average local clustering coefficient is given by:

$$ALCC = \frac{1}{N}\sum_{i=1}^{N} C_i.$$

**Global Efficiency (GE):** Let $L_{ij} = \frac{1}{a_{ij}}$ denote the pairwise length between any two nodes $i$ and $j$ (smaller weights result in larger lengths, and if $a_{ij} = 0$ then the pairwise length is infinite). Let $W_{ij}$ denote the set of finite sequences, or walks, $w = (L_{ik_1}, \dots L_{k_m k_{m+1}}, \dots, L_{k_M j})$ from node $i$ to $j$, where if $M = 0$, then the walk is given by $\left(L_{ij}\right)$, if $M = 1$, then the walks are given by $(L_{ik_1}, L_{k_1 j})$, for $k_1 \in \{1, .., N\}\backslash\{i, j\}$, and so on. Define the function: $L(w) = \sum w$ for all $w \in W_{ij}$, which gives the length of each walk. Then the distance between $i$ and $j$ is given by $d_{ij} = L(\overline{w})$, for any $\overline{w} \in \{w \in W_{ij} | L(w^*) \geq L(w)\ \forall\ w^* \in W_{ij}\}$. In other words, $d_{ij}$ is given by the shortest path length between the two nodes. If $L(w) = \infty\ \forall\ w \in W_{ij}$, then $d_{ij} = \infty$ (node $i$ and $j$ are not in the same connected component). The global efficiency is given by:

$$GE = \frac{1}{N(N - 1)}\sum_{i=1}^{N}\sum_{j \neq i}^{N}\frac{1}{d_{ij}},$$

where if $d_{ij} = \infty$ then $\frac{1}{d_{ij}} = 0$.

Generally, the network markers DStd, GE and AWCC are correlated with MD. In order to correct for this effect, these markers were normalized by their average value calculated over 500 randomized versions of the original network.

| Stage (# of epochs) | Diagnosis | Subjects (Cont/Epi) | Sex ratio (F/M) | Median age (Cont/Epi) | Cont-Epi age diff. (p-value) |
|---|---|---|---|---|---|
| Wake (31) | Cont | 9 | 2 (6/3) | 9 | 0.84 |
| | Epi | 22 | 0.47 (7/15) | 9.5 | |
| N1 (27) | Cont | 7 | 2.5 (5/2) | 8 | 0.18 |
| | Epi | 20 | 0.43 (6/14) | 10 | |
| N2 (21) | Cont | 6 | 2 (4/2) | 7.5 | 0.46 |
| | Epi | 15 | 0.25 (3/12) | 9 | |

**Table S1:** Number of epochs in each stage for epilepsy diagnosis, sex ratio (female/male), and age distribution.

| Subject | Sex | Age | Epilepsy | SCQ | Conners | Wake | Sleep N1 | Sleep N2 |
|---|---|---|---|---|---|---|---|---|
| PCB11 | Male | 13 | Focal | 9 | 4 | Yes | Yes | Yes |
| PCB12 | Male | 9 | Rolandic | 6 | 12 | No | Yes | Yes |
| PCB14 | Male | 8 | Encephalopathy | 27 | 8 | Yes | Yes | Yes |
| PCB15 | Male | 7 | No Epilepsy | 2 | 3 | Yes | Yes | Yes |
| PCB16 | Male | 9 | Generalized | 25 | 17 | Yes | No | No |
| PCB17 | Female | 8 | No Epilepsy | 8 | 8 | Yes | Yes | Yes |
| PCB19 | Female | 12 | No Epilepsy | 24 | 19 | Yes | Yes | Yes |
| PCB20 | Female | 9 | No Epilepsy | 11 | 4 | Yes | No | No |
| PCB25 | Male | 14 | Focal | 21 | 18 | Yes | Yes | Yes |
| PCB27 | Female | 9 | No Epilepsy | 27 | 17 | Yes | Yes | Yes |
| PCB28 | Female | 7 | No Epilepsy | 4 | 9 | Yes | Yes | Yes |
| PCB29 | Male | 15 | Rolandic | 10 | 0 | Yes | Yes | Yes |
| PCB3 | Male | 4 | No Epilepsy | 9 | 15 | No | Yes | Yes |
| PCB30 | Male | 14 | Generalized | 9 | 10 | Yes | Yes | Yes |
| PCB31 | Male | 12 | Focal | 16 | 4 | Yes | Yes | No |
| PCB32 | Male | 9 | Focal | 8 | 11 | Yes | Yes | Yes |
| PCB34 | Male | 6 | Encephalopathy | 12 | 13 | Yes | Yes | Yes |
| PCB36 | Female | 10 | Focal | 3 | 0 | Yes | Yes | No |
| PCB38 | Female | 12 | Generalized | 10 | 12 | Yes | Yes | No |
| PCB40 | Male | 13 | No Epilepsy | 21 | 17 | Yes | No | No |
| PCB44 | Female | 11 | Generalized | 6 | 1 | Yes | Yes | No |
| PCB45 | Male | 15 | No Epilepsy | 0 | 0 | Yes | No | No |
| PCB46 | Male | 10 | Focal | 18 | 20 | No | Yes | Yes |
| PCB47 | Male | 11 | Generalized | 14 | 11 | Yes | Yes | No |
| PCB5 | Male | 12 | Focal | 11 | 7 | Yes | Yes | Yes |
| PCB50 | Female | 10 | No Epilepsy | 5 | 3 | Yes | Yes | No |
| PCB51 | Male | 11 | Focal | 9 | 2 | Yes | No | No |
| PCB52 | Male | 6 | Focal | 5 | 14 | Yes | Yes | Yes |
| PCB58 | Female | 7 | Focal | 3 | 11 | Yes | Yes | Yes |
| PCB61 | Male | 6 | Focal | 4 | 20 | Yes | Yes | Yes |
| PCW3 | Female | 7 | Generalized | 2 | 18 | Yes | Yes | Yes |
| PCW4 | Female | 5 | Generalized | 7 | 4 | Yes | No | Yes |
| RPB1 | Male | 8 | Rolandic | 10 | 0 | Yes | No | No |
| RPB6 | Female | 9 | Rolandic | 17 | 1 | Yes | Yes | No |

**Table S2:** Subjects metadata, indicating sex, age, epilepsy diagnosis, SCQ and Conners raw score, and for which stages a suitable clean EEG epoch has been identified (see Methods for details).

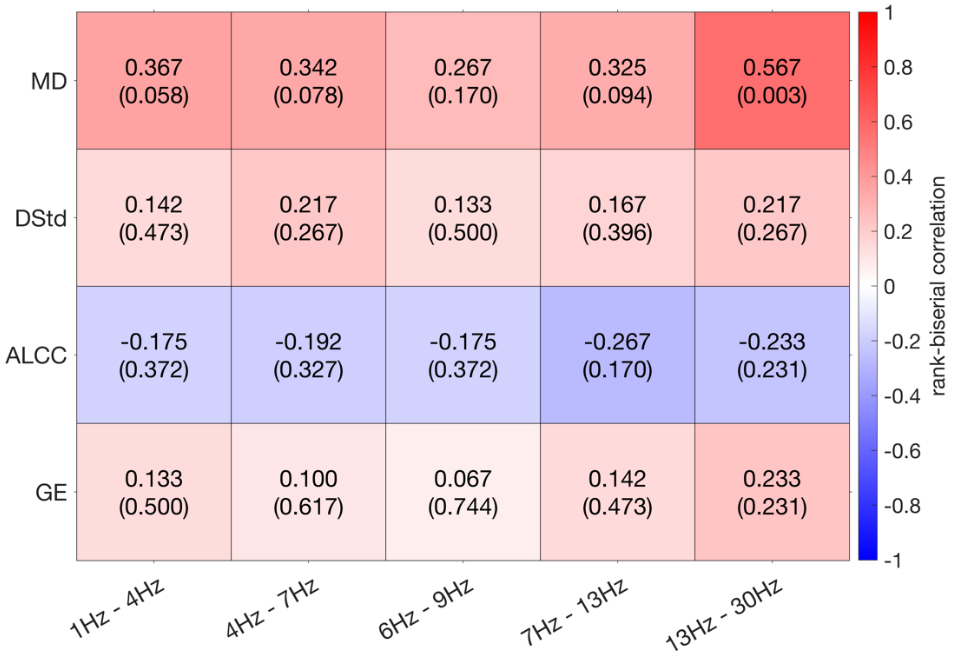


**Figure S1:** Rank-biserial correlation (and p-value) calculated for the comparison between Cont and Epi (a positive correlation means Cont > Epi) using the mean degree (MD), degree standard deviation (DStd), average weighted clustering coefficient (AWCC), and global efficiency (GE). These results were calculated using wake epochs.

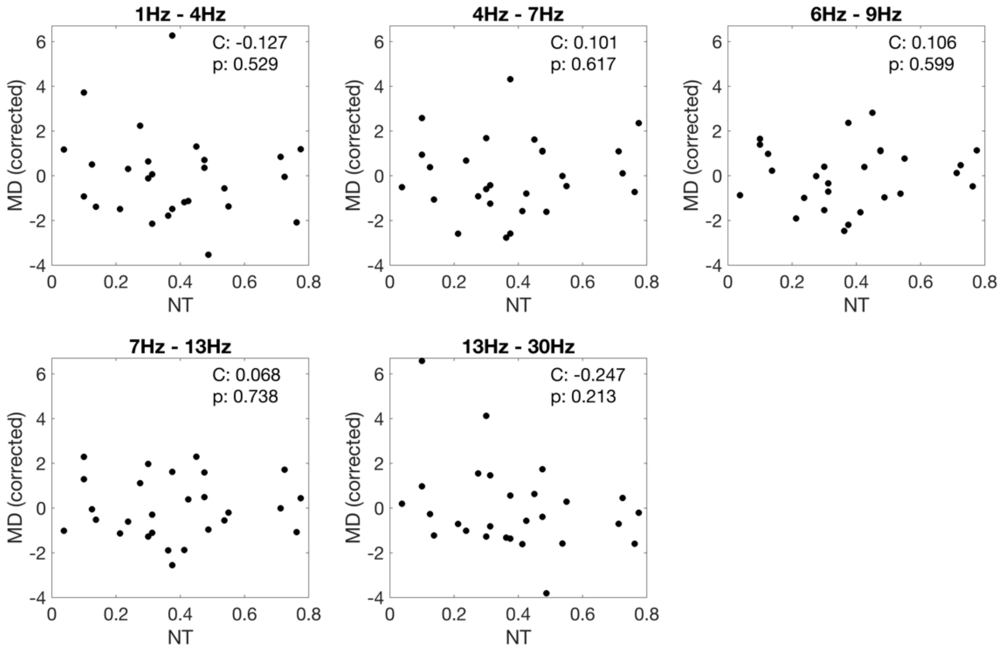


**Figure S2:** Spearman correlation and p-value (C and p) between neurodevelopmental trait index and mean degree (MD) corrected for age, sex, and epilepsy diagnosis. MD calculated using N1 epochs.

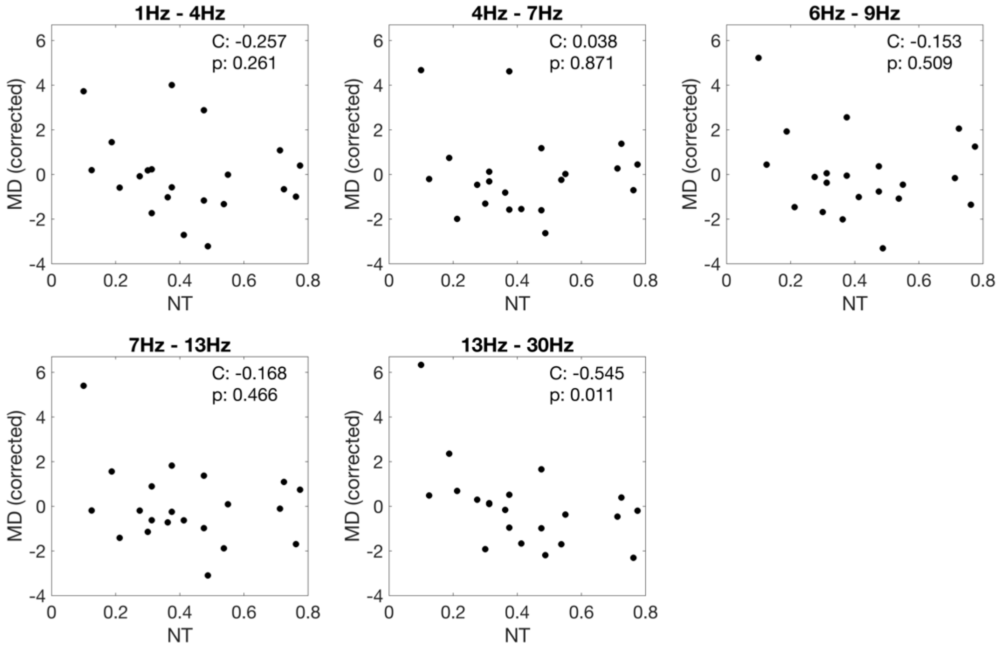


**Figure S3:** Spearman correlation and p-value (C and p) between neurodevelopmental trait index and mean degree (MD) corrected for age, sex, and epilepsy diagnosis. MD calculated using N2 epochs.

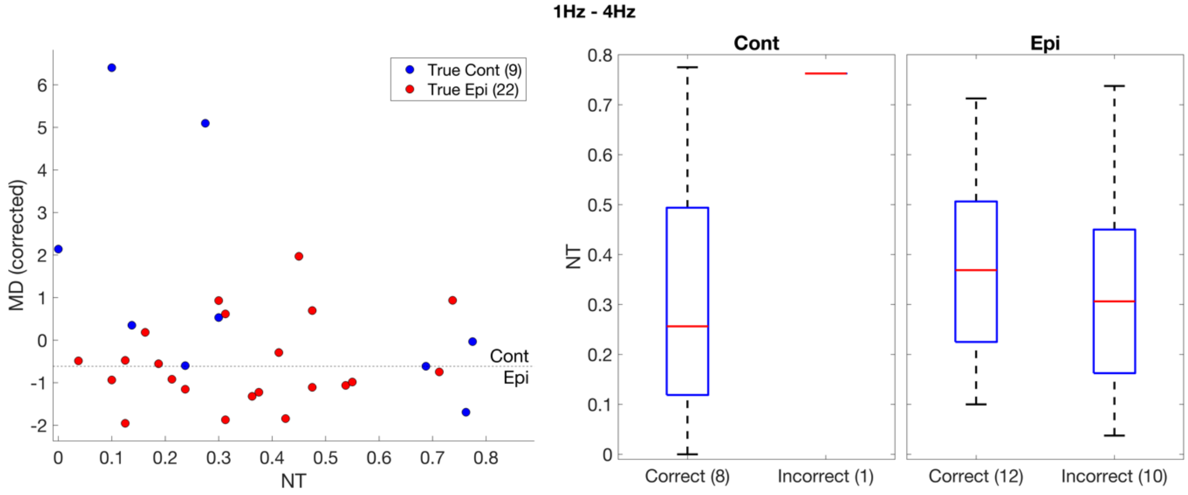


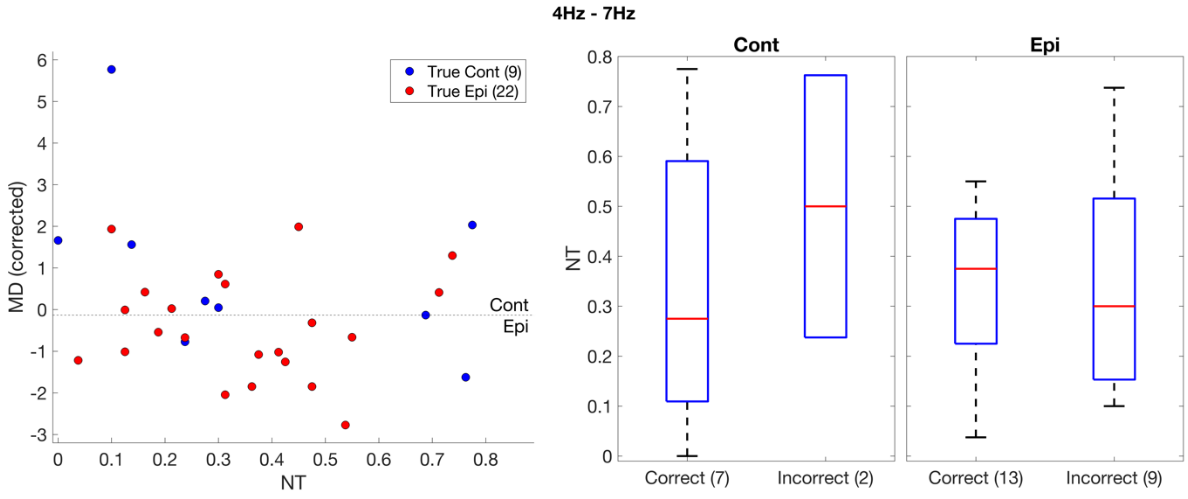

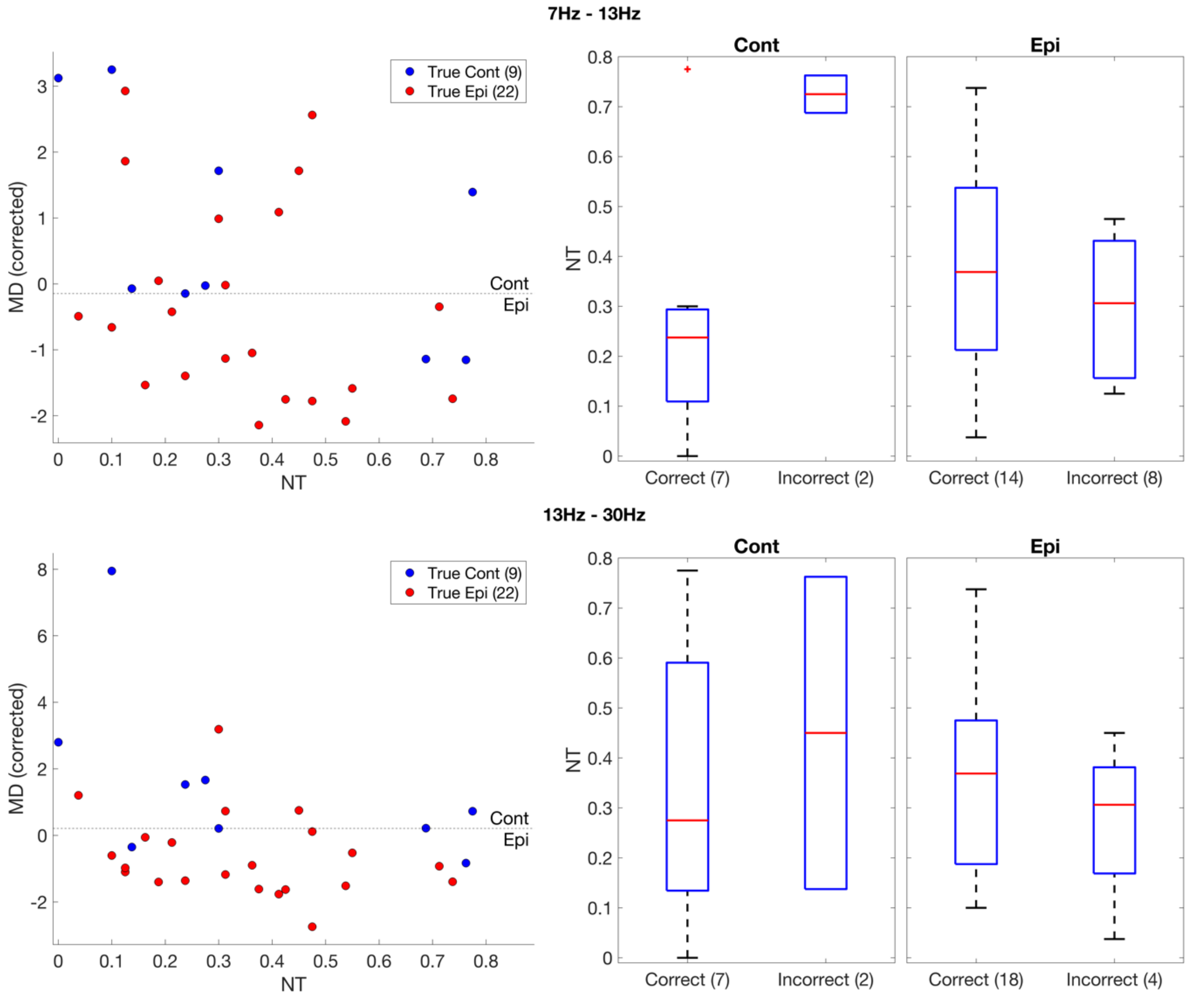


**Figure S4:** (Left) mean degree corrected for age, calculated using wake epochs, as a function of neurodevelopmental traits. Cont (Epi) are shown in blue (red). The dashed line represents the threshold of optimal balanced accuracy, for the separation between Cont and Epi. (Right) Comparison between neurodevelopmental traits of subjects classified correctly (Cont > threshold / Epi  < threshold) and incorrectly (Cont < threshold / Epi  > threshold). Each row of plots represents a different frequency band, indicated in the title.
